# iSeg: an efficient algorithm for segmentation of genomic and epigenomic data

**DOI:** 10.1101/184515

**Authors:** S.B. Girimurugan, Yuhang Liu, Pei-Yau Lung, Daniel L. Vera, Jonathan H. Dennis, Hank W. Bass, Jinfeng Zhang

## Abstract

**Background:** Identification of functional elements of a genome often requires dividing a sequence of measurements along a genome into segments where adjacent segments have different properties, such as different mean values. This problem is often called the segmentation problem in the field of genomics, and the change-point problem in other scientific disciplines. Despite dozens of algorithms developed to address this problem in genomics research, methods with improved accuracy and speed are still needed to effectively tackle both existing and emerging genomic and epigenomic segmentation problems.

**Results:** We designed an efficient algorithm, called iSeg, for segmentation of genomic and epigenomic profiles. iSeg first utilizes dynamic programming to identify candidate segments and test for significance. It then uses a novel data structure based on two coupled balanced binary trees to detect overlapping significant segments and update them simultaneously during searching and refinement stages. Refinement and merging of significant segments are performed at the end to generate the final set of segments. By using an objective function based on the *p*-values of the segments, the algorithm can serve as a general computational framework to be combined with different assumptions on the distributions of the data. As a general segmentation method, it can segment different types of genomic and epigenomic data, such as DNA copy number variation, nucleosome occupancy, nuclease sensitivity, and differential nuclease sensitivity data. Using simple t-tests to compute *p*-values across multiple datasets of different types, we evaluate iSeg using both simulated and experimental datasets and show that it performs satisfactorily when compared with some other popular methods, which often employ more sophisticated statistical models. Implemented in C++, iSeg is also very computationally efficient, well suited for large numbers of input profiles and data with very long sequences.

**Conclusions:** We have developed an effective and efficient general-purpose segmentation tool for sequential data and illustrated its use in segmentation of genomic and epigenomic profiles.

## Introduction

High throughput genomic assays, such as microarrays and next-generation sequencing, are powerful tools for studying genetic and epigenetic functional elements at a genome scale ^1^ A large number of approaches have been developed to exploit these technologies to identify and characterize the distribution of genomic and epigenomic features, such as nucleosome occupancy, chromatin accessibility, histone modifications, transcription-factor binding, replication timing, and DNA copy-number variations (CNVs). These approaches are often applied to multiple samples to identify differences in such features among different biological contexts. When detecting changes for such features, one needs to consider a very large number of segments that may undergo changes, and robustly calculating statistics for all possible segments is usually not feasible. As a result, heuristic algorithms are often needed to find the optimal solution for the objective function adopted by an approach. This problem is often called segmentation problem in the field of genomics, and change-point problem in other scientific disciplines.

Solving the segmentation problem typically involves dividing a sequence of measurements along the genome such that adjacent segments are different for a predefined criterion. For example, if segments without changes have a mean value of zero, then the goal could be to identify those segments of the genome whose means are significantly above or below zero. A large number of such methods have been developed for different types of genomic and epigenomic data ^2-15^. Many methods are designed for specific data types or structures, but it is challenging to find versatile programs for data with different properties and different underlying statistical assumptions. The previous methods fall into several categories including change-point detection ^2-4,8,11,14,16-23^, Hidden Markov models ^6,9,24-26^. Dynamic Bayesian Network (DBN) models ^27,28^signal smoothing ^29-32^, and variational models ^33,34^. For review and comprehensive comparison, please refer to ^13,35-38^.

Many currently available segmentation tools have poor performance, run slowly on large datasets, or are not straightforward to use. To address these challenges, we developed a general method for segmentation of sequence-indexed genome-wide data. Our method, called iSeg, is based on a simple formulation of the optimization problem^12^. Assuming the significance (i.e. p-value) of segments can be computed based on certain parametric or non-parametric models, iSeg identifies the most significant segments, those with smallest *p*-values. Once the segment with the smallest p-value is found, it will be removed from the dataset and segment with the second smallest p-value will be searched in the remaining of the data. The procedure repeats until no segments whose significance levels pass a predefined threshold. This simple objective function is intuitive from a biological perspective since the most statistically significant segments often biologically significant.

iSeg has several noteworthy features. First, the simple formulation allows it to serve as a general framework to be combined with different assumptions of underlying probability distributions of the data, such as Gaussian, Poisson, negative Binomial, or non-parametric models. As long as *p*-values can be calculated for the segments, the corresponding statistical model can be incorporated into the framework. Second, iSeg is a general segmentation method, able to deal with both positive and negative signals. Many of the existing methods, cannot deal with negative values in the data, because they are designed specifically for certain data types, such as genome-wide read densities, assuming data values with only zero or non-negative values. Negative values in genomic datasets can occur when analyzing data pair relationships, such as difference values or log-2 ratios commonly used with fold-change analysis. Currently, most methods segment profiles separately and compare the resulting segmentations. The drawback of such treatment is that peaks with different starting and ending positions from different segmentations cannot be conveniently compared, and peaks with different magnitudes may not always be distinguished. Taking differences from two profiles to generate a single profile overcomes these drawbacks. Third, to deal with cases where segments are statistically significant, but the biological significance may be weak, we apply biological significance threshold to allow practitioners the flexibility to incorporate their domain knowledge when calling “significant” segments. Fourth, iSeg is implemented in C++ with careful design of data structures to minimize computational time to accommodate, for example, multiple or very long whole genome profiles. Fifth, iSeg is relatively easy to use with few parameters to be tuned by the users.

Here we describe the method in detail, followed by performance analysis using multiple data types including simulated, benchmark, and our own. The data types include DNA copy number variations (CNVs) for microarray-based comparative genomic hybridization (aCGH) data, copy number variations from next generation sequencing (NGS) data, and nucleosome occupancy data from NGS. From these tests, we found that iSeg performs at least comparably with popular, contemporary methods.

## Materials and Methods

### Problem formulation

We adopt a formulation of the segmentation problem from previous methods^12^. The goal is to find segments with statistical significance higher than a predefined level measured by *p*-values under a certain probability distribution assumption. The priority is given to segments with higher significance, meaning that the segment with highest significance will be identified first, followed by the one with the second highest significance, and so on. We implemented the method using Gaussian-based tests (i.e. t-test and z-test), as used in other existing methods ^3,12,39^. Our method achieved satisfactory performance for both microarray and next generation sequencing data without modifying the hypothesis test. A more common formulation of the change-point problems is given in ^40^.

Consider a sample consisting of *N* measurements along the genome in a sequential order, *X_1_, X_2_,…, X_N_*, and

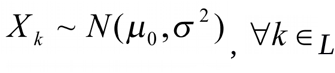

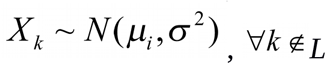

for some set of locations *L* (i.e. background, regions with no changes, etc.). The common assumption is that there are *M* non-overlapping segments with mean *^μ_1_, μ_2_,…,μ_M_^*, where *μ_i_ # μ_0_* and the union of these segments will form the complement of the set *L*. If the background level,*^μ_0_^*, is non-zero, the null hypothesis can be the coưesponding non-zero means. According to this model, it is possible for multiple segments with means different from *^μ_0_^* to be adjacent to each other. In addition, all the measurements are assumed to be independent. This assumption has been employed in many existing methods ^21,39^. A summary of existing methods that use such an i.i.d assumption and its properties are discussed in ^41^. The goal of a segmentation method is to detect all the *M* segments with means different from ^*μ_0_*^.

To illustrate, Figure 1(A) shows segments generated from Normal distributions with nonzero means where the rest of the data is generated from a standard Normal distribution. There are two computational challenges associated with the approach we are taking that also manifest in many previous methods. First, the number of segments that are examined is very large. Second, the overlaps among significant segments need to be detected so that the significance of the overlapping segments can be adjusted accordingly. To deal with the first challenge, we applied dynamic programming combined with exponentially increased segment scales to speed up the scanning of a large sequence of data points. To deal with the second challenge, we designed an algorithm coupling two balanced binary trees to quickly detect overlaps and update the list of most significant segments. Segment refinement and merging allow iSeg to detect segments of arbitrary length.

**Figure 1.**
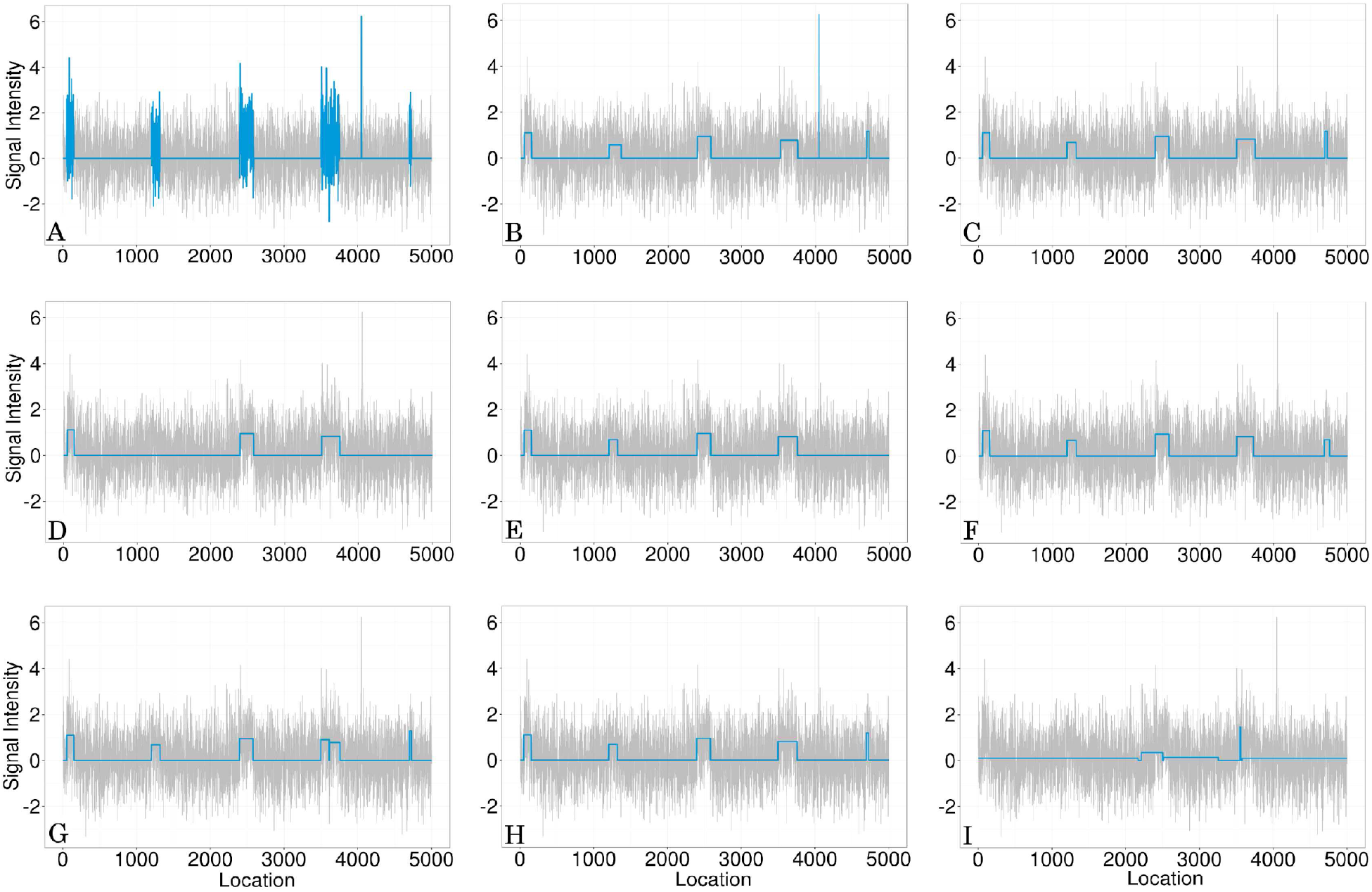
One of the simulated profiles and its detected segments obtained using iSeg. (A) The actual data with background noise and predefined segments. The segments with non-zero means are normally distributed with unit variance and means 0.72, 0.83, 0.76, 0.9, 0.7, and 0.6 respectively. The profiles shown here are normalized for an approximate signal to noise ratio of 1.0. The segments detected by iSeg (B) and other existing methods: snapCGH (C), mBPCR (D), cghseg (E), cghFLasso (F), HMMSeg (G), DNAcopy (H) and fastseg (I).

#### Computing p-values using dynamic programming

iSeg scans a large number of segments starting with a minimum window length, *W_min_*, and up to a maximum window length, *W_max_*. The minimum and maximum window lengths have default values of 1 and 300, respectively. This window length increases by a fixed multiplicative factor, called power factor (*ρ*), with every iteration. For example, the shortest window length is *W_min_*, and the next shortest window length would be *ρW_min_*. The default value for *ρ* is 1.1. When scanning with a particular window length, *W*, we use overlapping windows with a space of *W*/5. When ‘*W*’ isn’t a multiple of 5, numerical rounding (**ceil**) is applied. The aforementioned parameters can be changed by a user. We found the default parameters work robustly for all the datasets we have worked with. The algorithm computes *p*-values for candidate segments and detects a set of non-overlapping segments most significant among all possible segments.

Given the normality assumption, a standard test for mean is the one-sample student’s t-test, which is commonly found among many existing methods. The test statistic for this test is,

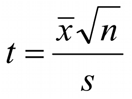

where 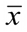 is the sample mean, s is the sample standard deviation, and n is the sample size. A drawback of this statistic is that it cannot evaluate segments of length 1. This may be the reason that some of the previous methods are not good at detecting segments of length 1. Although we can derive a test statistic separately for segments of length 1, the two statistics may not be consistent. To solve this issue, we first estimate the sample standard deviation using median absolute deviation and assume that the standard deviation is known. This allows us to use *z* statistic instead of *t* statistic and the significance of single points can be evaluated based on the same model assumption as longer segments. To calculate sample means for all segments to be considered for significance, the number of operations required by a brute force approach is ‘*C_b_’*.

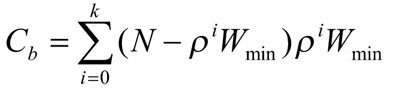

where, *ρ^k^ W_min_* ≤ *W_max_* and *ρ^k+1^W_min_* > *W_max_*.

Computation of these parameters (means and standard deviations) for larger segments can be made more efficiently by using the means computed for shorter segments. For example, the running sum of a shorter segment of length ‘m’ is given by,

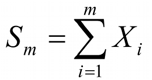

If this sum is retained, the running sum of a longer segment of length *r* (*r* > *m*) in the next iteration can be obtained as,

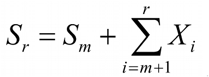

and the means for all the segments can be computed using these running sums. Now, the total number of operations *(C_b_^*^)* is

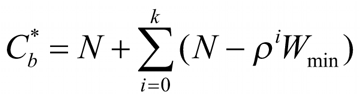

which is much smaller in practice than the number of operations (*C_b_*) without using dynamic programming. Computation of standard deviations is sped up using a similar process.

#### Detecting overlapping segments and updating significant segments using coupled balanced binary trees

When the p-values of all the segments are computed, we rank the segments by their *p*-values from the smallest to the largest. All the segments with *p*-values smaller than a threshold value, *p_s_*, are kept in a balanced binary tree (BBT1). The default value of *p_s_* is set as 0.001. Assuming a significance level (*α*) of 0.1, 100 simultaneous tests will maintain a family-wise error rate (FWER) bounded by 0.001 with Bonferroni and Sidak corrections. Thus, the cut-off is an acceptable upper bound for multiple testing. It can be changed by a user if necessary. The procedure for overlapping segment detection is described below as a pseudo-code. The set BBT1 stores all significant segments passing the initial significance level cutoff (default value 0.001). The second balanced binary tree (BBT2) stores the boundaries for significant segments. After the procedure, SS contains all the detected significant segments. The selection of segments using balanced binary tree makes sure that segments with small *p*-values will be kept, while those overlapping ones with bigger p-values will be removed.

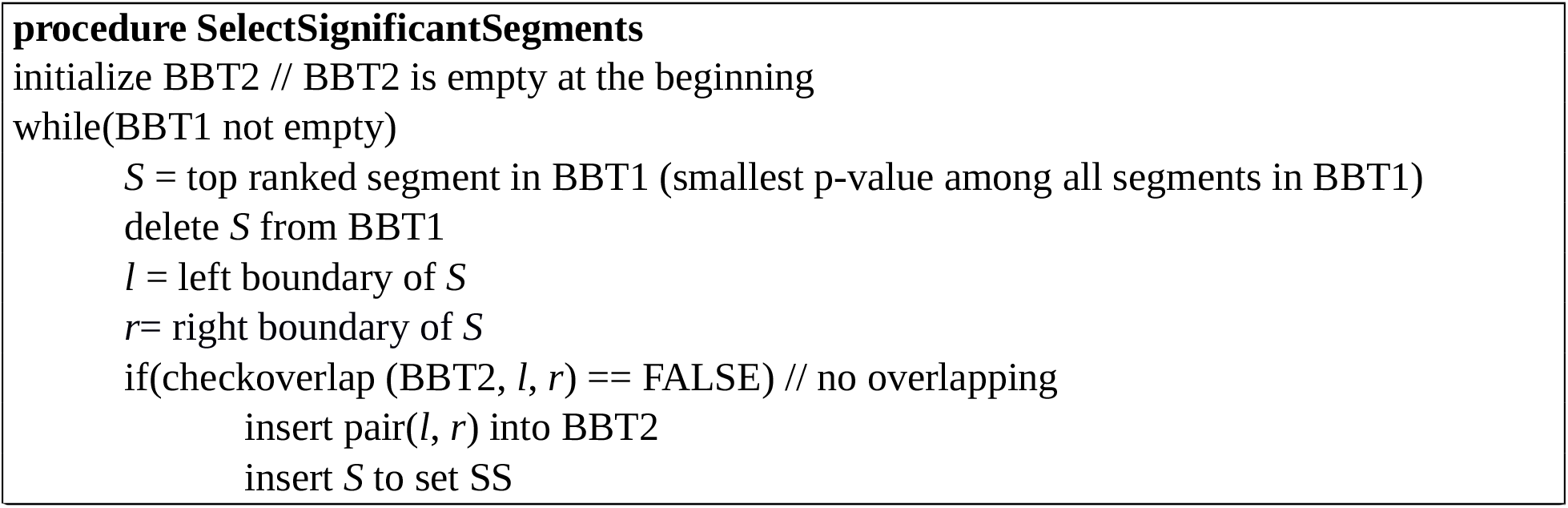

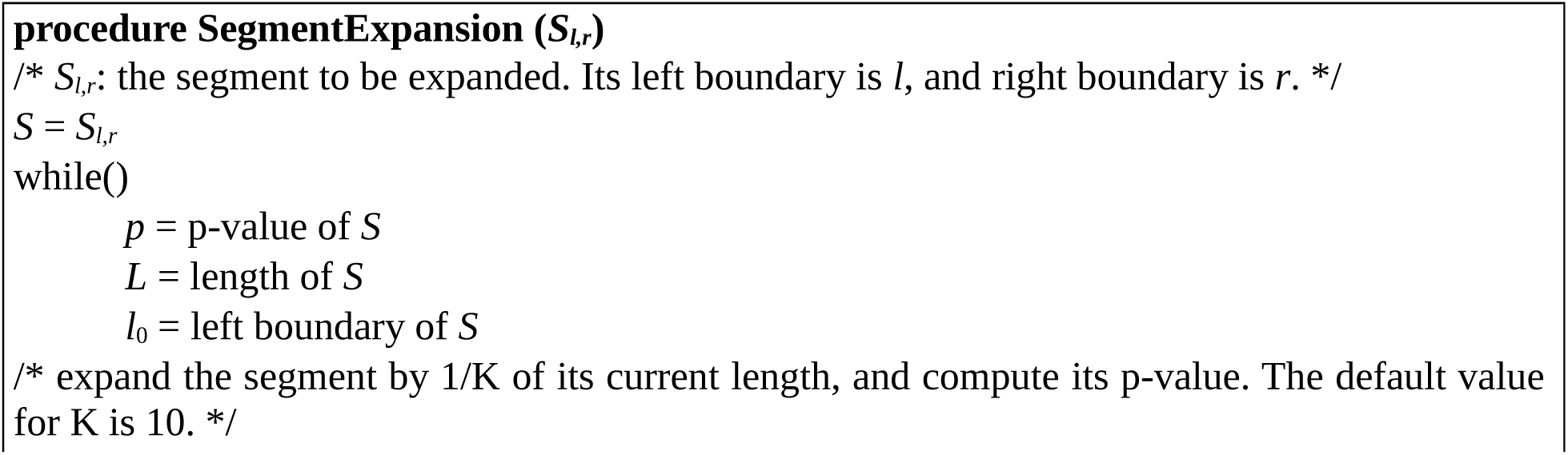
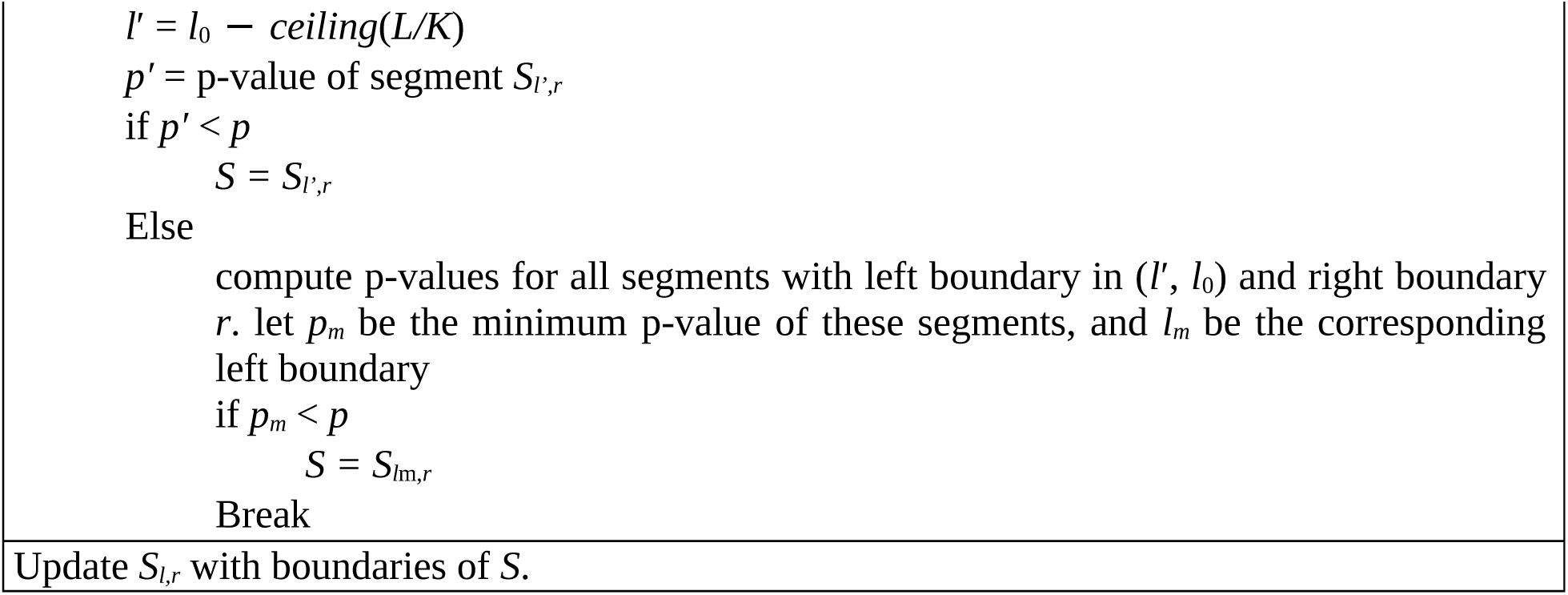

#### Refinement of significant segments

The significant segments are refined further by expansion and shrinkage. Without loss of generality, in the procedure (see SegmentExpansion text box) we describe expansion on left side of a segment only. Expansion on the right side and shrinkage are done similarly. When performing said expansion and shrinkage, a condition to check for overlapping segments is applied so the algorithm results in only disjoint segments.

#### Merging adjacent significant segments

When all the significant non-overlapping segments are detected and refined in the previous steps, iSeg performs a final merging step to merge adjacent segments (no other significant segments in between). The procedure is straightforward. We check each pair of adjacent segments. If the merged segment, whose range is defined by the left boundary of the first segment and the right boundary of the second segments, has a p-value smaller than those of individual segments, then we merge the two segments. The new segment will then be tested for merging with its adjacent segments iteratively. The procedure continues until no segments can be merged. With refinement and merging, iSeg can detect segments of arbitrary length— long and short. We added an option to merge only segments whose distances are no more than certain threshold, where distances are measured by the difference of the ending position of the first segment and the starting position of the second segment.

#### Multiple comparisons

In iSeg, p-values for potentially significant segments are calculated. Using a common p-value cutoff, for example 0.05, to determine significant segments can suffer from a large number of false positives due to multiple comparisons. To cope with the multiple comparisons issue, which can be very serious when the sequence of measurements is long, we use a false discovery rate (FDR) control. Specifically, we employ the Benjamini-Hochberg (B-H) procedure ^42^ to obtain a cutoff value for a predefined false discovery rate (α), which has a default value of 0.01, and can also be set by a user. Other types of cutoff values can be used to select significant segments, such as a fixed number of most significant segments.

#### Biological cutoff

Often in practice, biologists prefer to call signals above a certain threshold. For example, in gene expression analysis, a minimum of two-fold change may be applied to call differentially expressed genes. Here we add a parameter, *bc*, which can be tuned by a user to allow more flexible and accurate calling of significant segments. The default output gives four *bc* cutoffs: 1.0,1.5, 2.0 and 3.0.

#### Processing of the raw NGS data from Maize

Raw fastq files were clipped of 3' illumina adapters with cutadapt 1.9.1. Reads were, aligned to B73 AGPv3 (Schnable et al., 2009) with bowtie2 v2.2.8 (Langmead et. al, 2009), alignments with a quality <20 were removed, and fragment intervals were generated with bedtools (v2.25) bamtobed (Quinlan and Hall 2010). Fragments were optionally subset based on their size. Read counts in 20-bp nonoverlapping windows across the entire genome were calculated with bedtools genomecov and normalized for sequencing depth (to fragments-per-million, FPM). Difference profiles were calculated by subtracting heavy FPM from light FPM. Quantile normalization was performed using the average score distributions within a given combination of digestion level (or difference) and fragment size class.

## Results

We compare our method with several previous methods for which we were able to obtain executable programs: HMMSeg^6^, CGHSeg^14^, DNAcopy^4,22^, fastseg^3^ cghFLasso^32^, BioHMM-snapCGH^24^, mBPCR^11^, SICER^43^, PePr^44^ and MACS^45^. Among them, CGHSeg, DNAcopy, BioHMM-snapCGH, mBPCR and cghFLasso are specifically designed for DNA copy number variation data; MACS, SICER and PePr are designed for ChiP-seq data; and HMMSeg is a general method for segmentation of genomic data. Each method has some parameters that can be tuned by a user to achieve better performance. In our comparative study, we carefully selected parameters on the basis of the recommendations provided by the authors of the methods. For each method including iSeg, a single set of parameters is used for all data sets except where specified. Post-processing is required by some of the methods to identify significant segments.

In our analysis, performance is measured using F1-scores^46^ for all methods. F_1_-scores are considered as a robust measure for classifiers because they account for both precision and recall in their measurement. The F_1_-score is defined as,

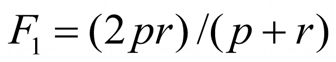

where *p* is precision and *r* is recall for a classifier. In terms of the true (TP) and false (FP) positives,

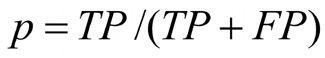

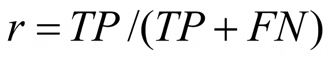

The methods CGHSeg, DNAcopy, and fastseg depend on random seeds given by a user (or at runtime automatically), and the F_1_-score at different runs are very similar but not the same. These methods were run using three different random seeds. The averages of the F_1_-score were used to measure their performance.

### Performance on simulated data

The simulated profiles were generated under varying noise conditions, with signal to noise ratios (SNR) of 0.5, 1.0 and 2.0, which correspond to poor, realistic and best case scenarios, respectively. Ten different profiles of length 5000 were simulated.

For each profile, five different segments of varying lengths were predefined at different locations. Data points outside of these segments were generated from normal distribution with mean zero. The five segments were simulated with non-zero means and varying amplitudes, or ease of detection, in order to assess the robustness of the methods. Because this set of simulated data resembles more of the DNA copy-number variation data, we used it to compare iSeg to methods designed for DNA copy number data. Figure 1 shows an example of the simulated data and the segments identified by iSeg and other existing methods. Figure 2(A) shows the performance of iSeg and other methods on simulated data with SNR = 1.0. We can see that iSeg, DNACopy and CGHSeg perform similarly well, with HMMseg and CGHFLasso performing a little worse while fastseg did not perform as well as the other methods. iSeg is also tested using a set of 10 longer simulated profiles, each with length 100000. Seven segments are introduced at varying locations along the profiles. iSeg still performs quite well in these very long profiles. The performance of these methods on long sequences is shown in Figure 2(B).

**Figure 2.**
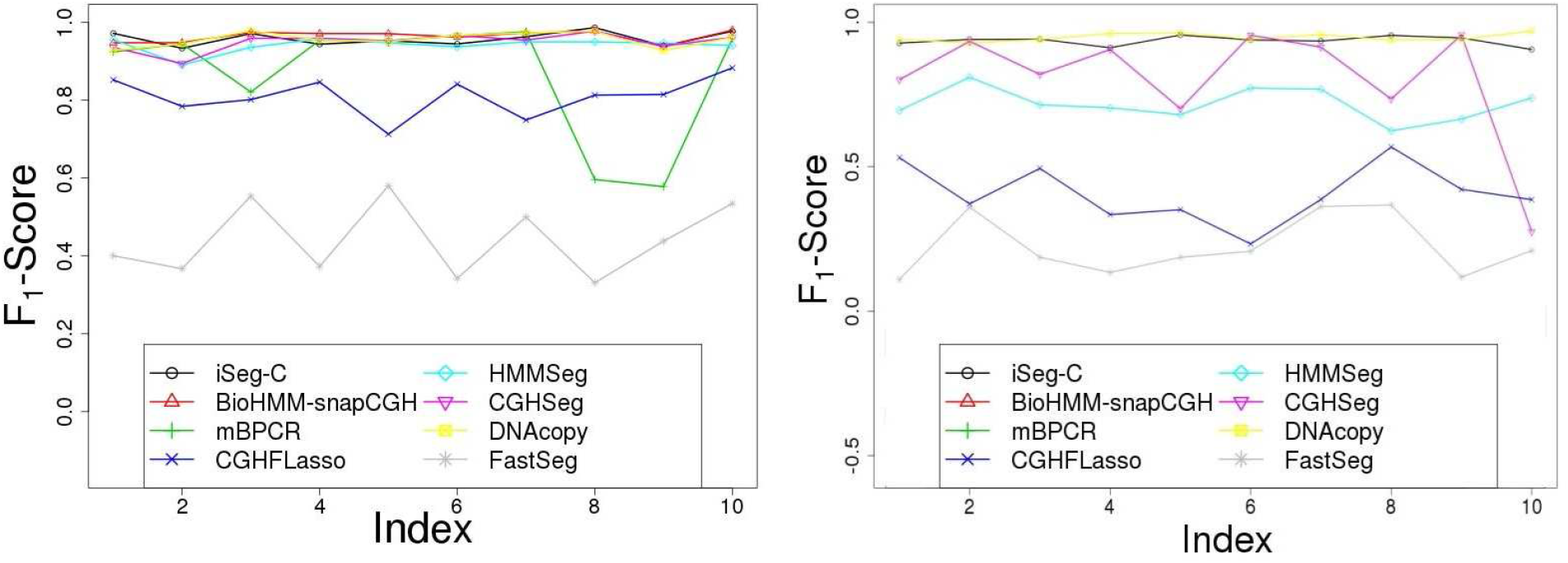
Comparison of F_1_-scores of various methods in analyzing different types of profiles. A. Simulated Profiles; B. Simulated Long Profiles (n=100K).

### Performance on experimental data

#### DNA copy-number variation (CNV) data

To assess the performance of iSeg on experimental data, we use three different datasets. They were the Coriell dataset^47^ with 11 profiles, the BACarray dataset^48^ with three profiles, and the dataset from The Cancer Genome Atlas (TCGA) with two profiles. The 11 profiles in Coriell datasets correspond to 11 cell lines: GM03563, GM05296, GM01750, GM03134, GM13330, GM01535, GM07081, GM13031, GM01524, S0034 and S1514. We constructed “gold standard” annotations using a consensus approach. We first ran all the methods using several different parameter settings for each method. The resulting segments were collectively evaluated using the test statistic described in the method. The set of gold standard segments were obtained using Benjamini-Hochberg procedure to account for multiple comparisons. The annotations derived using the consensus approach are provided as Supplementary material.

The 11 profiles from the Coriell dataset were segmented using iSeg and the other methods. Segmentation result for one of the profiles is shown in Figure 3, and the F_1_-scores are shown in Figure 4(A). The performance of iSeg is robust with accuracy above 0.75 for all the profiles from this dataset and it was found to be comparable to or better than other methods. For HMMSeg, both no-smoothing and smoothing were used. The best smoothing scale for HMMSeg was found to be 2 for the Coriell dataset. In Figure 3, we found that iSeg identified most of the segments. DNAcopy, fastseg, HMMSeg and cghseg missed single-point peaks, whereas cghFLasso, mBPCR and snapCGH missed some of the longer segments. The segmentation results for other profiles in Coriell dataset can also be found in the Supplementary material. We generated annotations using the consensus method for BACarray dataset similar to the Coriell dataset. The comparison of segmentation results for one profile of the BACarray dataset is shown in Figure 5, and the comparison of F1-scores is shown in Figure 4(B). iSeg returned better F1-scores than the other methods, consistent with the conclusions based on visual inspection.

**Figure 3.**
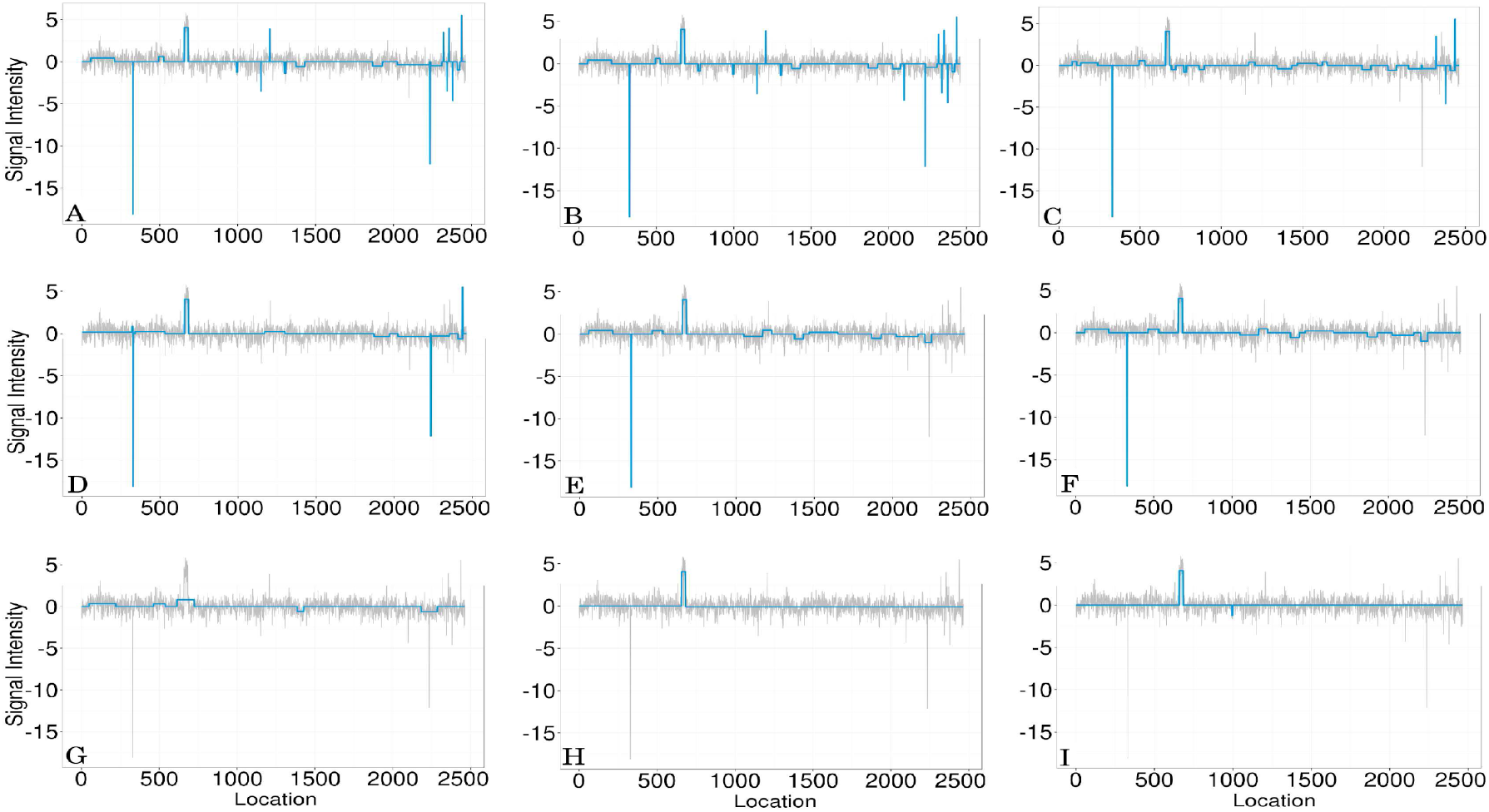
Comparison of segmentation results for one of the Coriell datasets. (A) The gold standard segmentation obtained using a consensus approach. Segmentation results of iSeg (B) and other existing methods: snapCGH (C), mBPCR (D), cghseg (E), cghFLasso (F), HMMSeg (G), DNAcopy (H) and fastseg (I).

**Figure 4.**
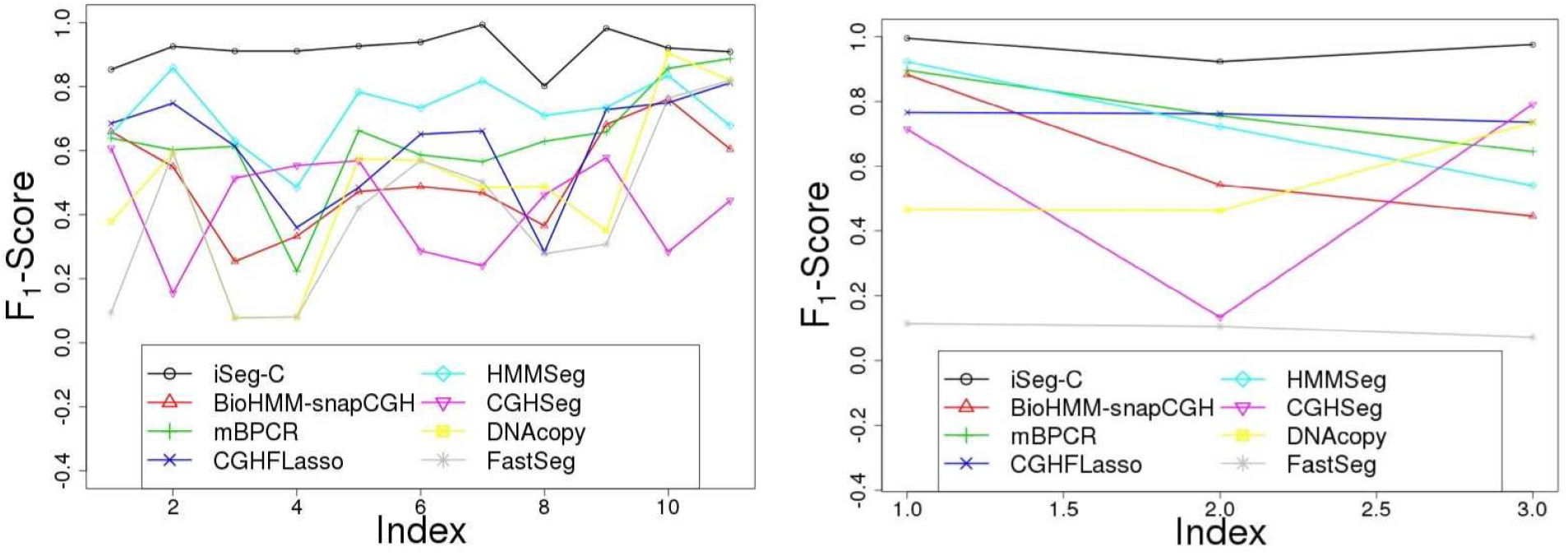
Comparison of F_1_-scores of various methods in analyzing different types of profiles. A. Coriell (Snijders et al.) profiles; B. BACarray profiles.

**Figure 5.**
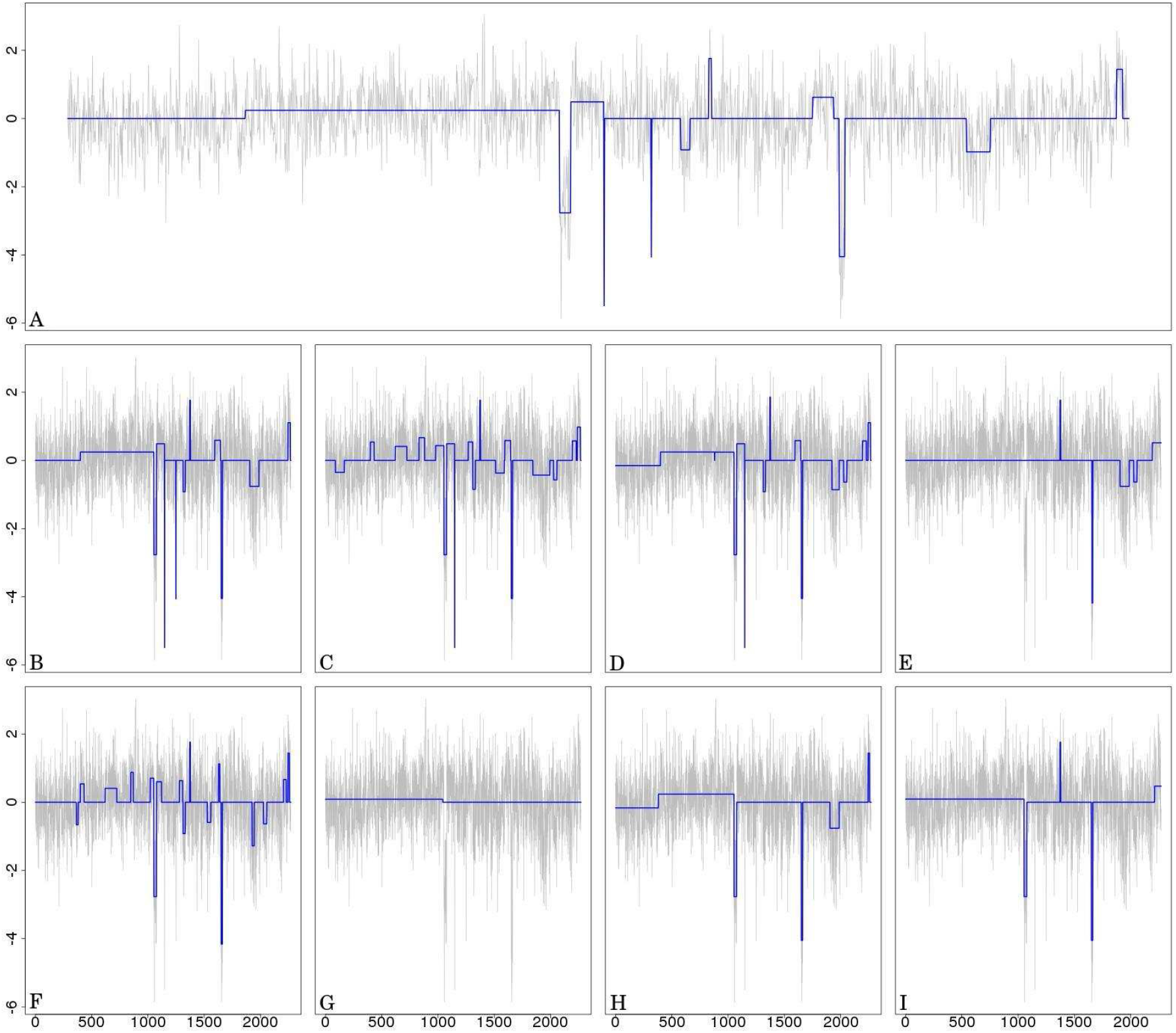
Comparison of segmentations for BACarray dataset. (A) The gold standard segmentation obtained using a consensus approach. Segmentation results of iSeg (B) and other existing methods: snapCGH (C), mBPCR (D), cghseg (E), cghFLasso (F), HMMSeg (G), DNAcopy (H) and fastseg (I).

For the TCGA datasets, since the profiles are rather long, we did not generate annotations using the consensus approach. Instead, we applied some of the methods to this dataset and compared their segmentation results visually (Figure 6). Again, we found that iSeg identified most of the significant peaks. In this test, DNACopy performed well overall, but tended to miss some of the single-point peaks, whereas other methods performed even less well.

**Figure 6.**
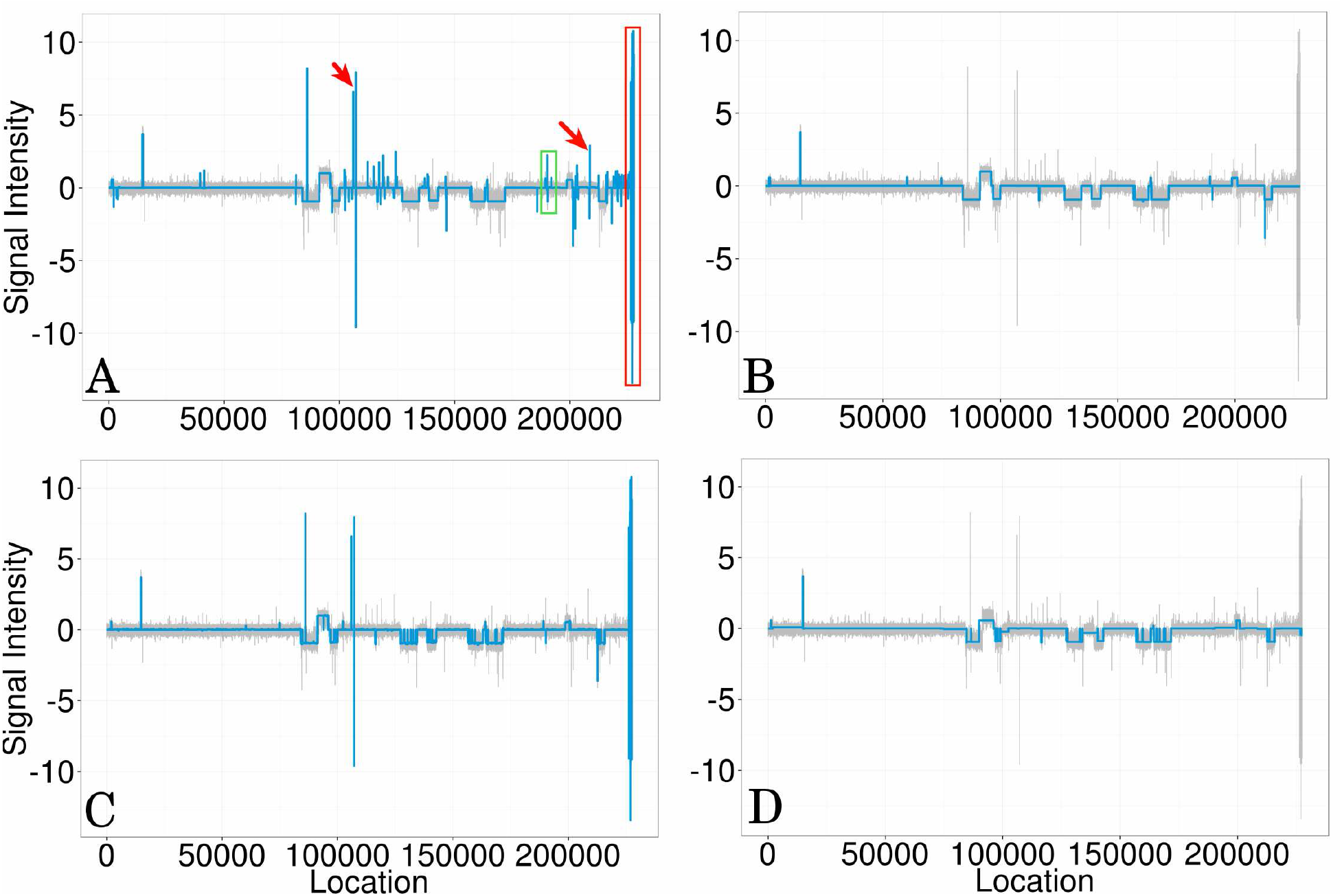
Comparison of segmentations for the TCGA dataset. The patient profile ID is TCGA-02-0007 and the data is supplied by the Harvard Medical School 244 Array CGH experiment (HMS). Segmentation results of iSeg (A) and other existing methods: DNAcopy (B), cghFLasso (C), and cghseg (D). The peaks pointed by the arrows and the region labeled by the red, and green squares are identified by iSeg, but not all of them are detected by the other three methods. Overall, iSeg consistently identifies all the significant peaks. Other methods often miss peaks or regions which are more significant than those identified.

We compared the computational time of iSeg, shown in Table 1, with those of the other methods, and found that iSeg is the fastest for the three test datasets. Notably, iSeg took much less time than the other methods for very long profiles (length 100000). This speed is achieved in part through dynamic programming and a power factor that provides rapid initial scanning of the profiles. The long profiles contain similar amount of data points that are signals (as opposed to background or noise) as the shorter profiles. The time spent on dealing with potentially significant segments is roughly the same between the two types of profiles. As a result, the overall running time of iSeg for the long profiles did not increase as much as that for the other methods. In summary, we observed that iSeg ran faster than the other methods, especially for profiles with sparse signals.

**Table 1.** Comparison of computational times (in seconds) on simulated data and Coriell data. These are total times required to process 10 simulated and 11 Coriell profiles.

| Method | Simulation<br>(SNR≈1.0, n=5000) | Simulation<br>(SNR≈1.0,<br>n=100K) | Coriell |
| --- | --- | --- | --- |
| iSeg (C++) | 0.164 | 1.223 | 0.294 |
| DNAcopy (R) | 2.267 | 60.343 | 3.098 |
| Fastseg (R) | 0.647 | 48.139 | 0.630 |
| CGHSeg (R) | 54.480 | 157.626 | 24.36 |
| HMMSeg (Java) | 0.543 | 160.790 | 0.552 |

### Differential nuclease sensitivity profiling (DNS-seq) data

We then tested our method on next generation sequencing data, for which discrete probability distribution models have been used in most of the previous methods. The dataset profiles were genome-wide reads from light or heavy digests (with zero or positive values) or difference profiles (light minus heavy, with positive or negative values)^49^. The difference plots are also referred to as sensitivity or differential nuclease sensitivity (DNS) profiles^49^.

#### Segmentation of single nuclease sensitivity profiles

Figure 7 shows the significant segments (peaks) called by iSeg together with those of two other methods, MACS and SICER. Visual inspection revealed that iSeg successfully segmented clear peaks and their boundaries. The performance of iSeg was at least comparable to MACS and SICER. Follow-up analyses on the segmentation results also demonstrated its capability in identifying biological interesting functional regions ^49,50^. iSeg results with different biological significance cutoffs (BC) are displayed as genome browser tracks beside the input profile data to guide inspection of the segmentation results. The default is to output three BCs: 1.0, 2.0, and 3.0, which are sufficient for most applications.

**Figure 7.**
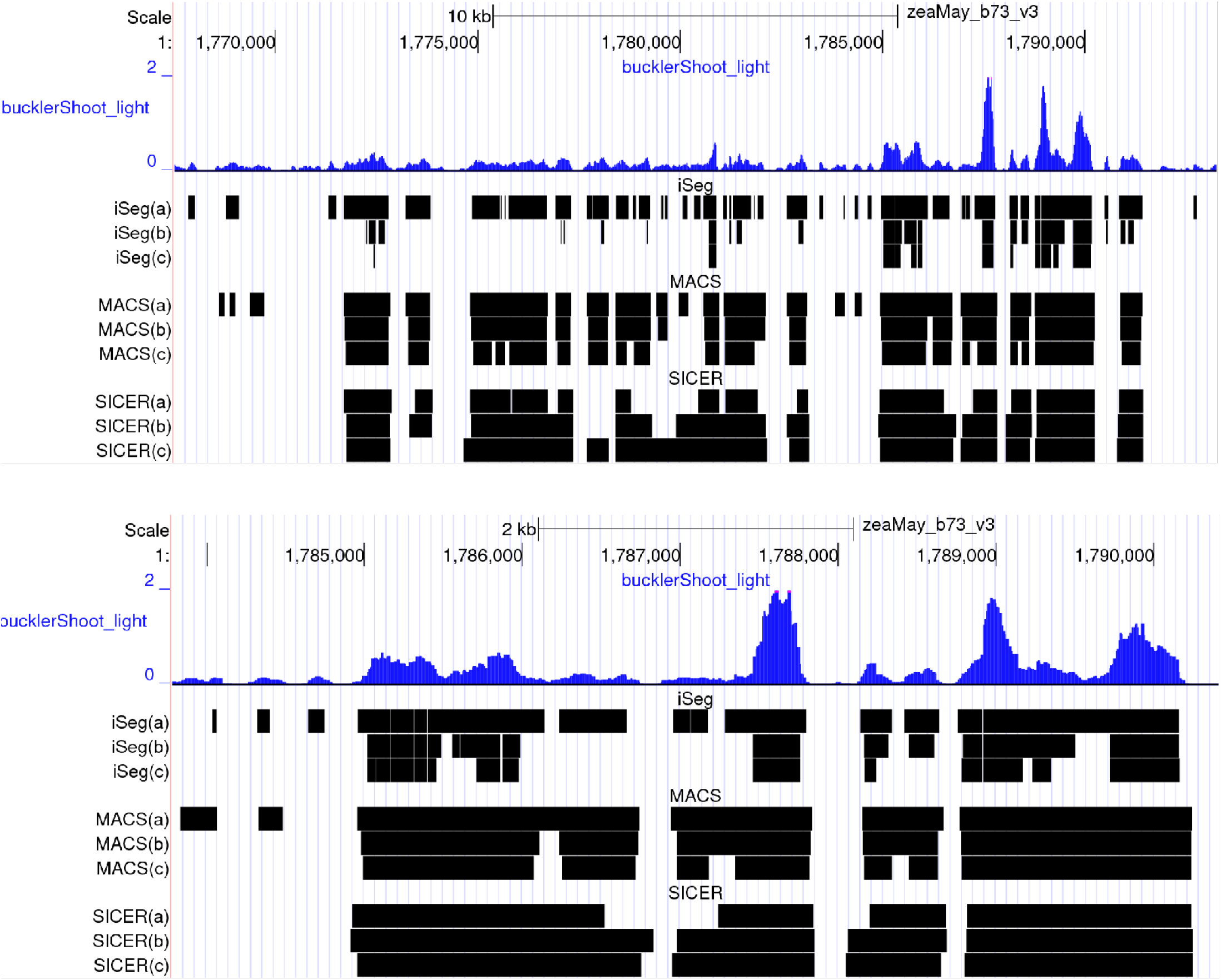
Demonstration of iSeg peak calling performance on Maize Epigenomics B73V3 nuprime **Buckler Shoot** CHIP-seq data. From top to bottom: First signal track is light group (**small fragment data only, color = blue**). Segment tracks are enriched regions called by: iSeg(a): iSeg calls with **bc = 1.0;** iSeg(b): iSeg calls with **bc = 2.0**; iSeg(c): iSeg calls with bc = 3.0; MACS(a): MACS calls on the light only group, **FDR=0.05; MACS(b)**: MACS calls on the light only group, FDR=0.01; MACS(c): MACS calls on the light only group, **FDR=0.001**; SICER(a): SICER calls on the light only group with **window size = 20 and gap size = 40;** SICER(b): SICER calls on the light only group with **window size = 30 and gap size = 60;** SICER(c): SICER calls on the light only group with **window size = 50 and gap size = 100;** All tracks are available at url

#### Segmentation of difference profiles with both positive and negative values

Difference profiles between two conditions can be generated by subtracting one profile from the other at each genomic location. Pairwise comparisons are of great interest in genomics, as they allow for tests of differences within replicates, or across treatments, tissues, or genotypes. Analyzing such profiles can preserve the range or magnitude of differences, adding power to detect subtle differences between two profiles, compared to approaches that rely on calling peaks in the two files separately. Figure 8 shows the segmentation of iSeg on a typical DNS-seq profile. When analyzing these sets of DNS data, we merge segments only when they are consecutive, meaning the gap between the two segments is zero. The length of gaps between adjacent segments that can be merged is a parameter tunable by users. iSeg successfully identified both positive (peak, positive peak) and negative (valley, negative peak) segments. Most existing ChiP-seq data analysis methods do not accommodate this type of data as input. To run MACS and SICER, we assigned the light and heavy digestion read profiles as the treatment and control files, respectively. Since the true biological significant segments are unknown, we compared the methods through careful visual inspections by domain experts. We found that iSeg performed satisfactorily and select it as the method of choice for analyzing the data from our own labs.

**Figure 8.**
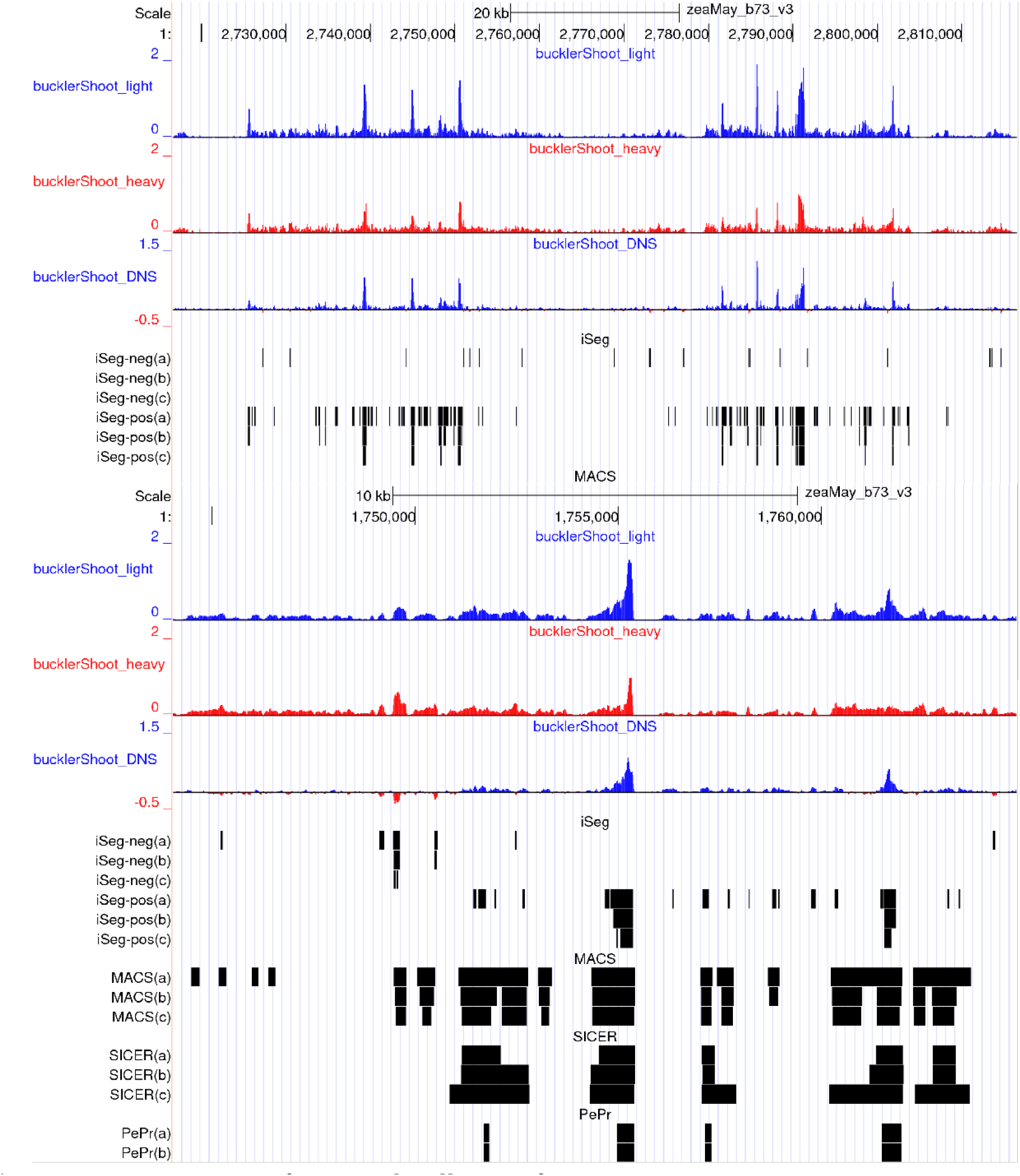
Demonstration of iSeg peak calling performance on Maize Epigenomics B73V3 nuprime Buckler Shoot CHIP-seq data. From top to bottom: First signal track is light group (treatment, color = blue); Second signal track is heavy group (control, color = red); Third signal track is difference track (light minus heavy, color code: positive = blue, negative = red).

## Conclusions

In this study, we designed an efficient method, iSeg, for the segmentation of large-scale genomic and epigenomic profiles. When compared with existing methods using both simulated and experimental data, iSeg showed comparable or improved accuracy and speed. iSeg performed equally well when tested on very long profiles, making it suitable for deployment in real-time, including online webservers able to handle large-scale genomic datasets.

In this study, we have assumed that the data follow a Gaussian (normal) distribution. The algorithm is not limited, however, to this distribution assumption. Other hypothesis tests, such as Poisson, negative binomial, and non-parametric tests, can be used to compute *p*-values for the segments. Data generated by next-generation sequencing (NGS) are often assumed to follow either a Poisson or a negative binomial distribution ^51,52^ However, a recent study showed that normality-based tests such as t-tests or Welch's t-test can perform equally well for NGS data ^53^, indicating Normal approximation of NGS data is likely adequate when analyzing NGS data, at least for some applications. For segmentation problems, when the segments tend to have relatively large sizes, the averages of signals within segments can be well approximated by normal distributions. This likely explains why iSeg performed well on NGS data. However, *p*-values of the segments can be computed based on either Poisson or negative binomial distributions, which can be directly used in the rest of the segmentation procedures. Such flexibility and general applicability make iSeg especially useful for segmenting a variety of genomic and epigenomic data.

The use of dynamic programming and a power factor makes iSeg computationally more efficient when analyzing very large data sets, especially with sparse signals as is typical for many types of NGS data sets. The refinement step identifies the exact boundaries of segments found by the scanning step. Merging allows iSeg to detect segments of any length. Together, these steps make iSeg an accurate and efficient method for segmentation of sequential data.

The statistic used in our method is very similar to that described for optimal sparse segment identification ^12^. However in that study, the segments are identified using an exhaustive approach, which will not be efficient for segmenting very large profiles. To speed up computation, the optimal sparse segment method ^12^ employs the assumption that the segments have relatively short length, which is not true for some datasets. In contrast, the algorithm designed in this study allowed us to detect segments of any length with demonstrably greater efficiency.

The gold standard generated using a consensus approach does not guarantee that the true optimal segments will be identified. In addition, the F_1_-scores may favor iSeg more as the test statistic used to generate the gold standard is not employed by the other methods. However, the statistic we used is based on model assumptions used by many existing methods. Visual inspection of segmentation results clearly shows how iSeg performs well in direct comparisons with other popular, contemporary methods. We expect that future research will benefit from the application of iSeg to compare multiple profiles simultaneously.

We have designed the method to make it flexible and versatile. This resulted in a number of parameters that users can tune. However, the default values work well for all the simulated and experimental datasets. In practice, to obtain satisfactory results, users are not expected to modify any parameters. The speed of iSeg would allow us and fellow researchers to implement it as an online tool to deliver segmentation results in real-time.

Segment tracks are enriched regions called by:

~~~
iSeg-neg(a): iSeg calls of negative regions on the difference track with **bc = 1.0;**
iSeg-neg(b): iSeg calls of negative regions on the difference track with **bc = 2.0;**
iSeg-neg(c): iSeg calls of negative regions on the difference track with **bc = 3.0;**
iSeg-pos(a): iSeg calls of positive regions on the difference track with **bc = 1.0;**
iSeg-pos(b): iSeg calls of positive regions on the difference track with **bc = 2.0;**
iSeg-pos(c): iSeg calls of positive regions on the difference track with **bc = 3.0;**
~~~

~~~
MACS(a): MACS calls with light as treatment and heavy as control, **FDR=0.05;**
MACS(b): MACS calls with light as treatment and heavy as control, **FDR=0.01;**
MACs(c): MACS calls with light as treatment and heavy as control, **FDR=0.001;**
~~~

~~~
SICER(a): SICER calls with light as treatment and heavy as control, **window size = 20 and gap size = 40;**
SICER(b): SICER calls with light as treatment and heavy as control, **window size = 30 and gap size = 60;**
SICER(c): SICER calls with light as treatment and heavy as control, **window size = 50 and gap size = 100;**
~~~

~~~
PePr(a): PePr calls with light as treatment and heavy as control, **broad peak** mode;
PePr(b): PePr calls with light as treatment and heavy as control, **sharp peak** mode;
~~~

## List of abbreviations

aCGH: (micro)array-based comparative genomic hybridization BBT: balanced binary tree;
B-H: Benjamini-Hochberg;
CNV: copy-number variation;
DBN: dynamic Bayesian Network;
FDR: false discovery rate;
FPM: fragment per million;
FWER: family-wise error rate;
MACS: model-based Analysis for ChIP-Seq;
NGS: next generation sequencing;
SNR: signal-to-noise ratio;
TCGA: The Cancer Genome Atlas.

## Supplementary materials

The zipped supplementary file submitted with this manuscript contains Coriell (Snijders et al.) profiles, BACarray profiles, and simulated data files.

## Declarations

### Ethics approval and consent to participate

Not applicable.

### Consent for publication

Not applicable.

## Availability of data and material

The data used in this study are available publicly or in the supplementary materials.

## Authors' contributions

JZ designed the core iSeg algorithm. JZ, HWB, DLV, and JHD were responsible for the study design and result interpretation. SBG, YL, and PYL performed data analyses. SBG wrote the first draft of the manuscript. All authors revised the manuscript and approved the final version.

## Acknowledgements

We would like to thank the researchers who have generated the data used in this study.

